# Discovery of *Drosophila melanogaster* from Wild African Environments and Genomic Insights into Species History

**DOI:** 10.1101/470765

**Authors:** Quentin D. Sprengelmeyer, Suzan Mansourian, Jeremy D. Lange, Daniel R. Matute, Brandon S. Cooper, Erling V. Jirle, Marcus C. Stensmyr, John E. Pool

## Abstract

A long-standing enigma concerns the geographic and ecological origins of the intensively studied vinegar fly, *Drosophila melanogaster*, a globally widespread species [1] which “has invariably appeared to be a strict human commensal” [2]. In spite of its sub-Saharan origins, this species has never been reported from undisturbed wilderness environments that might reflect its pre-commensal niche [3]. Here, we document the collection of 288 *D. melanogaster* individuals from African wilderness areas in Zambia, Zimbabwe, and Namibia. After sequencing the genomes of 17 flies collected from Kafue National Park, Zambia, we found reduced genetic diversity relative to town populations, elevated chromosomal inversion frequencies, and strong differences at specific genes including known insecticide targets. Combining these new genomes with prior data enabled us to gain novel insights into the history of this species’ geographic expansion. Our demographic estimates indicated that an expansion from southern Africa began approximately 10,000 years ago, with a Saharan crossing soon after, but expansion from the Middle East into Europe did not begin until roughly 1,400 years ago. This improved model of demographic history will provide a critical resource for future evolutionary and genomic studies of this key model organism. Our results add historical context to the species’ human association, and the opportunity to study wilderness populations opens the door for future studies on the biological basis of its adaptation to human environments.

## RESULTS AND DISCUSSION

### Drosophila melanogaster *persists in African wilderness*

*D. melanogaster* is among the most intensively-studied species in the world, and yet it remains “among the commonest species whose exact place of origin and even ancestry have never been satisfactorily explained” [2]. Its relatives are distributed across sub-Saharan Africa and nearby islands, and a biogeographic analysis proposed an ancestral range in western and central Africa [3]. Despite considerable efforts to collect *D. melanogaster* from this equatorial region, it was never discovered in undisturbed wilderness, instead occurring only in human-settled areas and “seminatural habitats” [3]. However, a recent population genomic analysis suggested that *D. melanogaster* originated in southern Africa: populations from Zambia and Zimbabwe have the species’ highest levels of genetic variation, whereas other populations may have lost diversity due to founder event bottlenecks during geographic expansion [4]. These findings raise the possibility that *D. melanogaster* originated (and might still persist) in wild environments of southern-central Africa, which are primarily characterized by seasonally dry Miombo and Mopane woodlands [5]. Although *D. melanogaster* has occasionally been sampled from human settlements near natural areas in Zimbabwe [6,7], its hypothesized persistence in wild Miombo/ Mopane forests [4] remains unconfirmed.

Here, we report the collection of *D.* Matobo National Park, the revelation of this species’ *melanogaster* in five distinct natural areas of Zambia, strong affinity for the local marula fruit [8] enabled the Zimbabwe, and Namibia (**Figure 1A-F**). These collection of 255 wilderness individuals. locations represent a gradient of remoteness from The collection of *D. melanogaster* from wild human habitation, ranging from 1.4 km to >55 km Miombo and Mopane woodlands is consistent from the nearest camp or village (**Table S1**). At with its hypothesized pre-commensal origin from these environments [4]. These woodlands feature important seasonal fluctuations in fruit resources [9], temperature, and especially precipitation (**Figure 1G-I**). An origin from such variable environments (as opposed to humid equatorial forest) might contribute to the species’ more robust thermal, desiccation, and starvation tolerances than related species [10,11].

**Figure 1.**
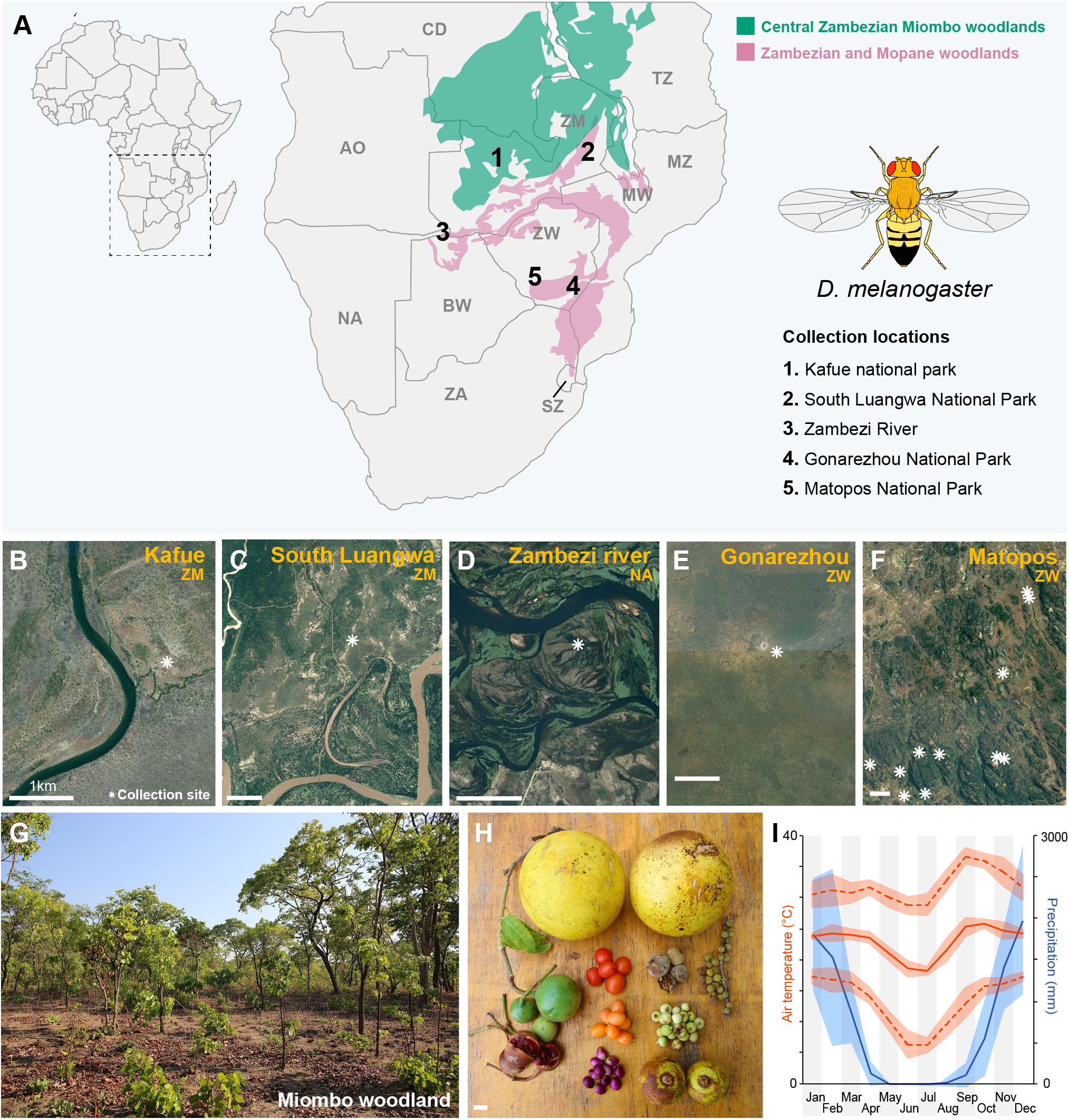
Collection environment and genomic differentiation of the Kafue population of *D. melanogaster.* (**A**) Locations of five D. melanogaster wilderness collections. (**B-F**) Satellite imagery of these sites. (**G**) Miombo woodland environment. (**H**) A sample of the diverse array of fruits present in this environment. (**D**) Fruits left on the ground by foraging monkeys offer possible fly breeding sites. (**E**) Climate graph for Livingstone, Zambia, depicting an extended dry season and a wide range of temperatures. See also **Table S1**.

### *Wild* D. melanogaster *have distinct patterns of genomic diversity*

In Kafue National Park (Zambia) we sampled a single male in Miombo wilderness over 4 km from any human structure, while multiple females and males were caught within ~4 km of an intermittently-used camp site (**Table S1**). To begin to understand the biology of wilderness *D. melanogaster* and its relationship to commensal populations, we sequenced individual genomes of 5 females and 12 males, including the more remote male individual [12] (STAR Methods).

Based on sequence comparisons [13,14] against town-sampled genomes with previously-inferred karyotypes [15], the Kafue population displayed some of the highest chromosomal inversion frequencies ever observed in *D. melanogaster.* The overall proportion of inverted chromosome arms was 2.3 times higher in Kafue than in the Siavonga, Zambia town population (*Table S2; z* = 6.49; *P* < 0.0001), and higher than the previous *D. melanogaster* collections scored by cytological [16] or genomic [15,17] methods depicted in **Figure 2**.

**Figure 2.**
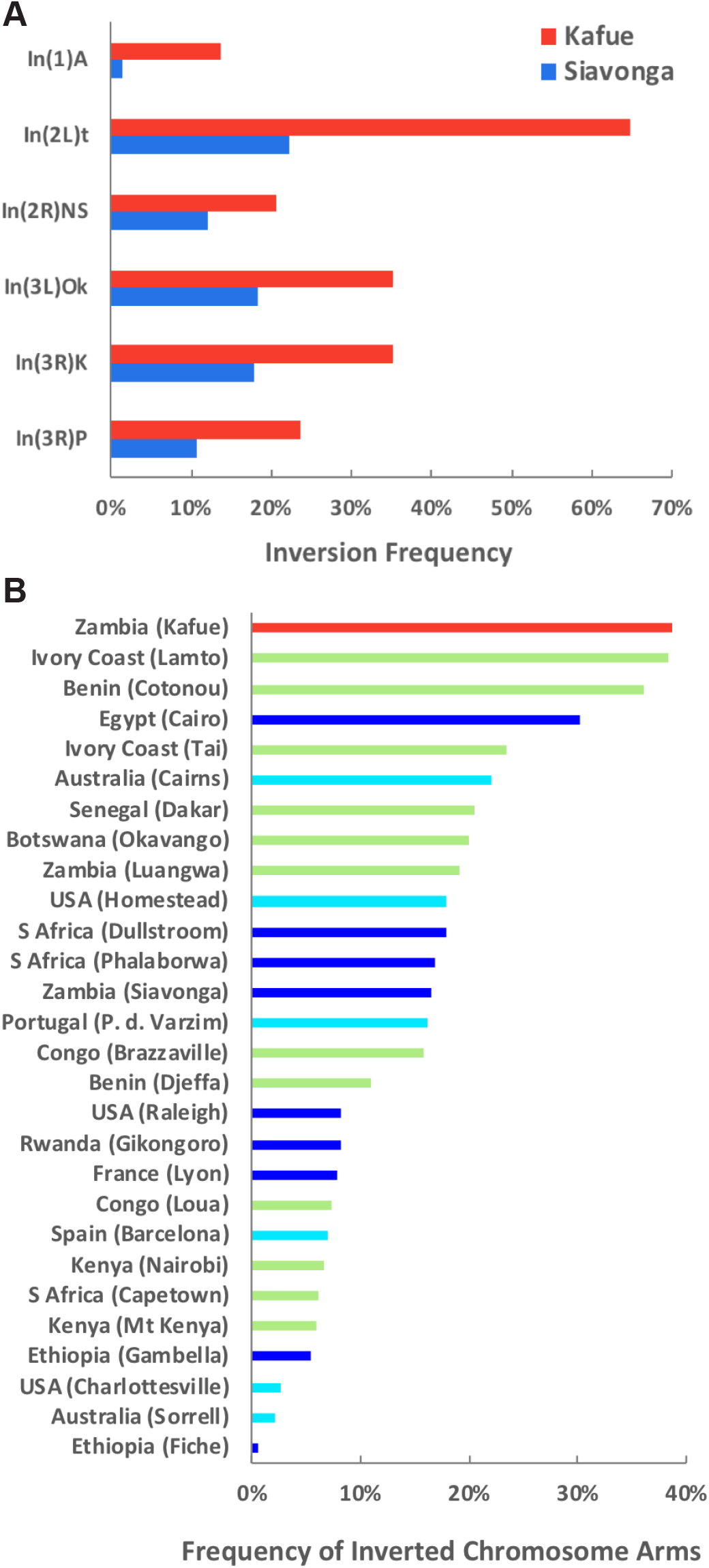
Elevated inversion frequencies in the Kafue population. (**A**) For all six chromosomal inversions common in this region, the Kafue park population yielded higher rearrangement frequencies than the Siavonga town population. (**B**) Kafue had a higher proportion of inverted chromosome arms (across X/2L/2R/3L/3R) than previous D. *melanogaster* collections from Africa or elsewhere. Inversions were computationally inferred from individual genomes (dark blue) [11] or pooled genomic samples (light blue) [16], or scored cytologically (green) [15]. Frequencies were compiled for populations with sample sizes of at least 20, and for the six inversions in (**A**), plus In(1)Be, In(3L)P, and In(3R)Mo, which are not common in Zambia. Among pooled samples [16], only the populations yielding the highest and lowest inversion frequencies from each continent are depicted. See also **Table S2**.

At least three distinct hypotheses could account for this preponderance of inversions. First, high inversion frequencies could be an ancestral characteristic, with frequencies declining during the adaptation and geographic expansion of commensal populations (analogous to the decline of inversion frequencies as the species has adapted to colder environments [18,19]). Second, high inversion frequencies could represent a derived characteristic, with local inversion haplotypes creating coadapted gene complexes [20] that link wilderness-adapted alleles in a park population to buffer against migration from larger town populations. Third, if the Kafue population represents a relatively small local deme (as suggested below), in which inbreeding may frequently expose recessive lethal/deleterious alleles [21], then inversion heterozygotes might avoid such fitness costs in light of their lower levels of genomic homozygosity. Further experiments and analysis will be needed to clarify the evolutionary forces shaping inversion frequencies in park and town populations.

Principle Components Analysis revealed overall genetic similarity between Kafue flies and southern African town populations, in line with a modest F_ST_ of 0.028 between Kafue and Siavonga (**Figure S1**). The more remote wilderness-collected individual clustered by PCA with other Kafue genomes), consistent with their membership in the same local population. Surprisingly, the Kafue genomes largely held a subset of the variation contained in the Siavonga, Zambia town population. Nucleotide diversity was reduced by 17% in the park sample (0.731% versus 0.876%; permutation P < 0.00001), which could not be explained by differences in haploid versus diploid genotype calling (STAR Methods).

The lower diversity of the Kafue genomes does not align with the predictions of a domestication bottleneck in the history of our commensal populations, although further genomic analysis of wilderness populations will be needed to assess the generality of diversity relationships. Instead, the reduced diversity of Kafue could hint that this local park population is part of a metapopulation in which individual demes have lower and potentially fluctuating population sizes that reduce local genomic diversity. However, this pattern does not exclude the possibility of the Kafue population being reintroduced from a Siavonga-like commensal source population, and experiencing a founder event or subsequent bottleneck. Thus, we pursued demographic inference in order to clarify the history of divergence and migration among park and town populations.

### *The History of* D. melanogaster

The availability of population genomic data from park and town populations offers the potential to illuminate the history of this species within and beyond its ancestral range. We therefore used fastsimcoal [22], a frequency-based demographic inference method, to estimate historical parameters relating the Kafue sample and previously sequenced town samples [15]. We began with the town populations, since many of them have larger sample sizes than Kafue and their somewhat greater differentiation may yield clearer estimates of divergence times, before returning to the question of the park population’s history. We deployed an iterative model-building approach to estimate the species’ expansion history. After starting with two populations, we added one additional population at a time, fixing previously estimated parameters not directly relevant to a given population’s addition. This strategy eliminated the need to estimate a prohibitive number of demographic parameters simultaneously. Since we could not investigate every possible population topology, we focused on the *D_xy_* distance tree obtained from genomic data [4], except for specific model comparisons noted below. We separately obtained results for chromosome arms X, 2R, and 3L (the arms with adequate samples of inversion-free genomes for most populations of interest). For brevity, we mainly focus below on median arm point estimates and their associated 95% confidence intervals for each parameter discussed below (STAR Methods), while full results are presented in **Table S3**.

Among the town populations, we estimated that Zambia and Rwanda populations of D. melanogaster split approximately 10,173 years ago (CI 8,690-13,146; **Figure 3**). This split may reflect the timing of *D. melanogaster* expanding beyond its southern-central African ancestral range. We estimated that the Cameroon-Oku sample split from the Rwanda lineage ~9,952 years ago (CI 8,606-12,822), which overlaps with the Zambia-Rwanda split, suggesting a relatively rapid onward expansion into west-central Africa. Our estimates indicate that our Ethiopian populations split off an estimated 7,556 years ago (CI 7,038-8,325). This sub-Saharan expansion would have occurred within a broad interval (7-16 kya) in which African environments were becoming warmer and wetter following the Last Glacial Maximum [23], potentially impacting both the distribution of wild habitat for *D. melanogaster* and human activity in the region. These sub-Saharan split times contrast with one study’s 72 kya divergence estimate between Zambia and a lumped panel of four western African populations [24]. However, they are similar to the ten pairwise sub-Saharan divergence times from another recent study [25]. Finally, we estimated a more recent divergence between our lowland and highland Ethiopian samples (based on only the X chromosome, which met sample size thresholds) at 2,069 years (CI 1,797-2,352), suggesting that occupation of the Ethiopian plateau was a more recent event.

**Figure 3.**
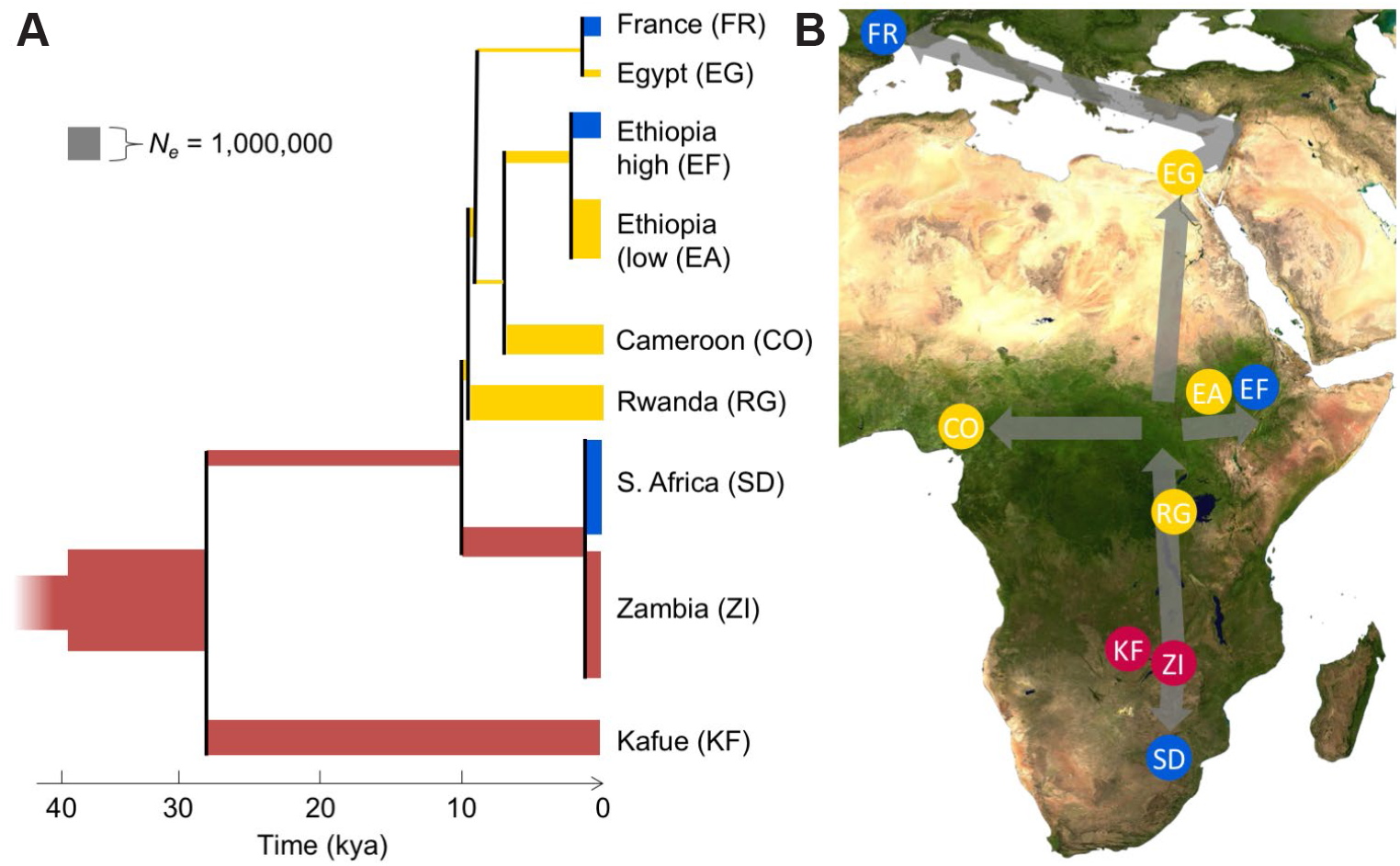
An estimated history of sampled *D. melanogaster* populations. (**A**) For the focal population tree identified from distance relationships and model likelihood comparisons, median arm point estimates are depicted for each estimated divergence time and population size. (**B**) The locations of the included populations are indicated. Colors of population branches (**A**) and circles (**B**) indicate populations and lineages within the apparent ancestral range (red), and those in regions occupied by roughly 10 kya (yellow) and by approximately 1-2 kya (blue). See also **Figure S1, Table S3, Table S4, and Table S5**.

Significant interest has centered on the timing of the species’ expansion out of sub-Saharan Africa, entailing a population bottleneck that created a major axis of genetic differentiation. Previous demographic studies [26-28] were limited to single African (Zimbabwe) and European (Netherlands) populations, and often assumed that the split between these populations occurred at the same time as the trans-Saharan founder event. In contrast, we investigated more geographically detailed models in which an Egypt/France branch split off from either the Cameroon, Ethiopia, or Rwanda lineages, or from internal branches between them. Of these, a split from the internal branch between the Rwanda and Cameroon lineages offered the highest likelihood for all three arms (**Table S4**), with an estimated split time of 9,793 years ago (CI 9,762-9,941). Hence, the species’ trans-Saharan expansion may have extended from its expansion across equatorial Africa, at a time when the Sahara was beginning to grow less arid [23]. There is evidence that fig cultivation in the Middle East began more than 11,000 years ago [29], which may have aided the persistence of commensal *D. melanogaster* outside of sub-Saharan Africa. Our estimated Saharan crossing ~10 kya is more recent than some previous population genetic estimates of ~16-23 kya [26-28]. This difference partly stems from our use of an empirically-based estimate of 15 generations per year [30], versus the previous conjecture of 10. Our estimates may also be aided by the inclusion of equatorial African populations, whereas most previous studies fixed the out-of-Africa bottleneck time to equal the divergence time between European and southern African populations.

We also report the first estimate of the expansion of *D. melanogaster* from the Middle East into Europe, at just 1,435 years ago based on arms X/3L (joint CI 1,192-1,517). This estimate offers context for the expansion of this species into cooler environments, although it may not have reached northern Europe until much later [1]. The European expansion may have been facilitated by historical factors such as Byzantine rule of the eastern Mediterranean from Constantinople (beginning in the 4th century CE) or Islamic control of Iberia (beginning in the 8th century CE), and by the concurrent expansion of grape and citrus cultivation in southern Europe [31,32].

Having addressed the history of range expansion, we returned to the Kafue sample. We explored four models in which Kafue branched from (1) the lineage ancestral to all town populations, (2) the shared Zambia-Siavonga and South Africa branch, (3) the Zambia-Siavonga terminal branch, or (4) the internal branch ancestral to Rwanda and the other non-southern populations. Of these, model 1, with Kafue as the sister to all other populations, had the best likelihood for two of three arms (**Table S4**), and so we focused on this topology. Model 4, which was supported by arm 2R, would still imply that the Kafue lineage had split from the others by roughly 10 kya. For model 1, we obtained an estimate of 29,106 years for the divergence between Kafue and the town populations, though with significant uncertainty in this date (CI 21,382-46,371). Although some temporal and topological uncertainty clearly remains, our estimated split time between geographically proximate Zambia populations is consistent with *D. melanogaster* having a deep history in this region, and suggests that the Kafue population was not recently reintroduced from a Siavonga-like town population.

We simultaneously added a northeastern South African population to our model, resulting in a divergence time estimate of 1,045 years from the Siavonga lineage (CI 782-1,297). A relatively recent expansion southward from the ancestral range could relate to Bantu groups’ arrival in this region beginning ~1,500 years ago and the accompanying onset of agriculture [33].

Because we focused on the chronology of range expansion and analyzed many populations, we implemented a simple model of population size, with two epochs in the ancestral population and one for each internal/terminal branch (**Figure 3**). Our estimates implied growth in the ancestral range, in line with the excess of rare alleles observed in sub-Saharan populations [12,15]. Natural selection may also contribute to such long-term growth signals [34,35], but the relatively recent parameters that differentiate our populations should be less affected [36]. The population sizes of other branches align with the progressive loss of genetic diversity during the species’ geographic expansion [4]. Whereas ancestral *N_e_* begins at ~1.65 million and reaches ~3.9 million in the Zambia lineage, the lowest value is roughly 100,000 for the Egypt/France lineage encompassing the trans-Saharan expansion (although the initial founder event size within that lineage was surely lower than our constant size estimate). Estimated migration rates ranged from roughly 10-9 to 10-5, with the highest values occurring between geographically proximate populations. (**Table S3**).

### Potential adaptive differences between park and town populations

The overall genetic similarity of park and town populations should improve our ability to detect allele frequency differences at specific genes that may result from local adaptation, including genetic changes related to the evolution of human commensalism. We therefore applied *Population Branch Excess* (*PBE*, a recently developed F_ST_-based statistic [37]) to identify genomic windows with unusually strong frequency differences between the Kafue population and the two southern African town populations mentioned above (Zambia-Siavonga and South Africa-Dullstroom).

Notably, the six strongest outlier regions included significantly higher *PBE* values than expected under neutrality from any window in the genome based on simulations using our demographic estimates (**Figure 4A; Table S6**; STAR Methods). The top window was centered on the transcription factor *lilliputian*, in which a 5’ UTR variant absent from 156 analyzed town genomes was present in 13 of 20 Kafue alleles. In light of its highly pleiotropic functions, which impact cell size, developmental patterning, and behavior, the specific role of this gene in local adaptation between park and town populations remains an open question. Aside from a low recombination region very close to the third chromosome centromere (which includes 36 genes), the strongest outlier regions also included two regions centered on poorly understood genes that both reach their highest expression level in the testes: the predicted phosphatase inhibitor *CG12620* and the predicted ubiquitination gene *CG31807.*

**Figure 4.**
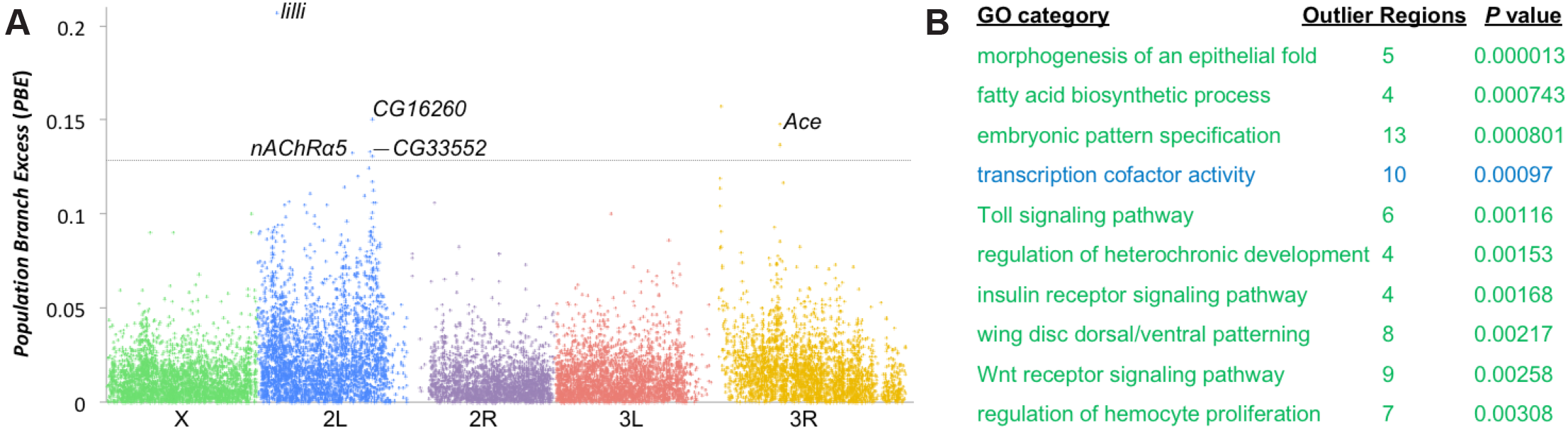
Potential genomic targets of adaptive differences between wild and town populations. (**A**) A plot of *Population Branch Excess* (*PBE*) indicates genomic windows (averaging 4 kb in length) that have elevated genetic differentiation between the Kafue population and town populations, with labeled examples described in the text. The dotted line indicates genome-wide significance based on neutral demographic simulations. (**B**) The gene ontology categories most enriched for *PBE* outliers are given, which include biological process (green) and molecular activity (blue) functions, after excluding redundant categories. See also **Table S6** and **Table S7**.

The other two top outliers both centered on neurotransmission-related genes that encode known insecticide targets: *Acetylcholine esterase* (*Ace*) and *nicotinic Acetylcholine Receptor α5* (*nAChRα5*). *nAChRα5* is part of the receptor targeted by neonicotinoids and some other insecticdes [38,39]. *Ace* is a major determinant of resistance to organophosphate and carbamate insecticides [40]. Known resistance mutations at Ace have been primarily reported among non-African populations [41]. Three such variants (at residues 161, 265, and 330) segregate among our Zambian genomes, with all showing lower frequencies in the park population (3% each in Kafue, versus 13%, 5%, and 5% in Siavonga). These modest differences are overshadowed by intronic SNPs that display frequency differences of ~50% between these park and town populations. It will therefore be of interest to test for additional resistance variants in African populations, and for differences in insecticide resistance between wilderness and commensal populations.

We then performed gene ontology (GO) enrichment analysis on *PBE* outliers to identify functional categories that may hold adaptive differences between the park and town populations. Some of the top GO categories pertained to developmental and gene regulatory functions that could have a wide range of potential impacts (**Figure 4B; Table S7**). Among the more specific processes implicated were insulin signaling and immune regulation categories (Toll signaling, hemocyte proliferation). Insulin signaling genes have shown strong clinal differentiation in North America [42]. Most immunity processes did not show elevated differentiation among five worldwide populations [43], but some immunity genes displayed parallel SNP frequency changes associated with the parallel adaptation to colder climates [19]. Although we cannot yet confirm which of these processes may have been targeted by local adaptation between park and town populations, results of this exploratory analysis provide hypotheses for phenotypes, genes, and processes that may underlie adaptation to wild versus anthropogenic environments.

### Conclusions

In this study, we have clarified an enduring conundrum, the origins and history of one of world’s most widely-studied organisms, *Drosophila melanogaster.* We have found that *D. melanogaster* continues to occupy wild environments in southern-central Africa. Alongside the concurrent discovery that *D. melanogaster* favors the indigenous marula over any other tested fruit [7], these results complement population genomic findings [4] suggesting that this region may represent the ancestral range of the species predating human commensalism. We have compared genomes from park and town samples to substantially clarify the history of this species’ geographic expansion: we have estimated the time-scale of its spread within sub-Saharan Africa, its Saharan crossing, and its spread into Eurasia, while connecting these events to relevant aspects of climate and human history.

We have also shown that a national park population from Zambia displays unique genetic features, including (1) reduced genetic diversity, (2) a surprisingly high frequency of polymorphic chromosomal inversions, and (3) elevated genetic differentiation in certain genes functional categories related to insecticide resistance, development, immune regulation, and insulin signaling. These patterns merit further study in other wild *D. melanogaster* populations. More broadly, the opportunity to study wilderness populations of *D. melanogaster* will add new dimensions to this evolutionary and genetic model system. Future studies may reveal behavioral, morphological, and physiological differences between wilderness and commensal populations.

Combining these phenotypic analyses with molecular studies, quantitative trait mapping, and population genetics may provide exceptional opportunities to uncover the genetic basis of an insect’s evolutionary transition to human commensalism. Insights obtained from this genetically tractable system may prove particularly relevant to the evolution of insect agricultural pests, as well as the adaptation of natural populations to novel environments.

## SUPPLEMENTAL INFORMATION

Supplemental Information includes Methods, one figure, and seven tables, and can be found with this article online.

## ACKNOWLEDGEMENTS

We are grateful for the kind hospitality and assistance we received from Chris and Charlotte McBride of McBride’s Safari Camp. We also thank Russ Corbett-Detig for advice on inversion calling and Nandita Garud for providing the locations of Ace SNPs. This work was funded by grants from the Crafoord Foundation and Vetenskapsrådet to MCS, by NIH grants to JEP (R01 GM111797), DRM (R01GM121750) and BSC (R35 GM124701), and by an NSF grant to DRM (1737752).

## AUTHOR CONTRIBUTIONS

MCS, SM, DRM, and BSC performed wilderness fly collections. EJ assisted with collections. QDS generated genome sequence data. QDS, JDL, and JEP performed population genomic analyses. JEP, QDS, MCS, DRM, and BSC wrote the manuscript.

## CONTACT FOR REAGENT AND RESOURCE SHARING

Further information and requests for resources should be directed to and will be fulfilled by the Lead Contact, John Pool.

## METHOD DETAILS

### Fly collection methods

We collected flies using standard traps, which were constructed from plastic bottles and baited with fruit (mango, banana, and local varieties) mixed with baker’s yeast. Traps were hung from trees at ~1.5 m above ground level and emptied after 24 h.

### Genomic sequence data collection

Genomic sequence data was collected from individual wild-caught, ethanol-preserved flies from the Zambia Kafue sample. These included 5 females (KF4, KF5, KF10, KF11, and KF20.) and 12 males (KF1, KF2, KF3, KF6, KF7, KF8, KF9, KF18, KF21, KF22, KF23, and KF24). Single fly DNA was isolated using the NEBNext^®^ Ultra^™^ DNA Library Prep Kit for Illumina (New England Biolabs E7370). Genomic sequencing was conducted using the Illumina HiSeq 2000 platform with 100 bp paired end reads, with ~300 bp inserts. Sequencing depth per genome ranged from 14X to 51X, with a median depth of 23X. Reference alignment and consensus sequence generation was as previously described^*12*^ for inbred strains. To facilitate the analysis of fully heterozygous chromosomes (*i.e.* all except male X chromosomes), we produced diploid consensus sequences with ambiguity codes representing heterozygous genotype calls. Assessing the outcomes of diploid versus haploid genotype calling, we found that female Kafue X chromosomes had lower nucleotide diversity than male Kafue X chromosomes (0.71% vs. 0.76%). However, X-linked diversity for the (haploid-called) Zambia-Siavonga sample was higher still (0.84%), confirming the lower genetic diversity of the Kafue population regardless of the ploidy of genotype calling. The generated Kafue files, along with published genomes^*15*^, were used for the calculation of genome-wide statistics based on genetic distances, and for the analyses described below.

## QUANTIFICATION AND STATISTICAL ANALYSIS

### Principle Components Analysis and Inversion Detection

To examine potential genetic differentiation between the Kafue sample and previously-studied town-collected populations from Southern Africa, we used the Principle Components Analysis (PCA) implementation of Patterson *et al*.^*13*^ (Eigensoft version 6.0.1). Whenever sequences of mixed ploidy were compared (heterozygous Kafue chromosomes versus haploid, homozygous, or hemizygous chromosomes from other populations), sites were categorized as X-linked with individuals labeled as female or male based on ploidy. Since inversions can alter genetic variation across whole chromosome arms^*4,14*^, we focused PCA on standard chromosome arms. For published individual strain genomes, inversion calls had previously been documented^*12*^. As previously shown^*14*^, inverted chromosome arms consistently have reduced genetic distance in breakpoint regions with respect to each other. For the Kafue genomes, suspected inversions were readily identified via examination of genetic distances around inversion breakpoints, in comparison with published standard and inverted chromosomes from southern Africa genomes^*12*^. These results were consistent with PCA-based groupings when inverted arms were included.

### Demographic Estimation

In order to better understand the expansion history of *D. melanogaster*, demographic parameters were estimated using fastsimcoal 2.6^*22*^. In addition to the Kafue genomes, we added data from other published *D. melanogaster* genomes from the Drosophila Genome Nexus^*12*^, including town-collected samples from Zambia (ZI), South Africa (SD), Rwanda (RG), Cameroon (CO), Ethiopia (lowland EA and highland EF), Egypt (EG), and France (FR). For each model investigated, we ran fastsimcoal for 100 independent runs, each with 100,000 coalescent simulations and 50 expectation-maximization cycles, and then selected the model with the smallest difference between observed and expected likelihoods to obtain point estimates.

Estimated parameters included population split times, symmetric migration rates between geographically adjacent populations, and population sizes (two epochs in the ancestral population, one size for each internal and terminal branch). Ranges of uniform priors are given in Table S3. Population size calculations were based our provided mutation rate of 6.6 × 10^-9 *S1*^. Conversion of generations (provided directly by fastsimcoal) to years was based on an estimate of 15 generations per year^*28*^.

Parameter estimates were separately obtained for chromosome arms X, 2R, and 3L. The unscaled output of fastsimcoal (raw generations, allelic migration probabilities) helps to make estimates from the X chromosome and autosomes more comparable. However, we might naturally expect lower population size estimates for the X, and we therefore rescale our reported X-linked *N_e_* estimates by a factor of 4/3, corresponding to the assumption of *N_f_* = *N_m_.* In general, we obtained similar results from X-linked and autosomal data (Table S3).

Because inversions may have distinct histories reflecting the action of natural selection and can affect genetic variation across whole arms^*4,14*^, we included only genomes that were inversion-free for a given chromosome arm. Higher inversion frequencies on arms 2L and 3R led us to omit them from our demographic analysis. To minimize the impact of direct selection on our analyzed sites, we included only short intron (bp 8 to 30 of introns ≤65 bp in length^*S2*^) and fourfold synonymous sites. To reduce the impact of linked selection, we only analyzed sites with regional sex-averaged recombination rates of at least 1 cM/Mb^*S3*^.

To obtain input data from each population for each analyzed chromosome arm, we chose sample sizes that jointly maximized this quantity along with the number of retained sites. We then excluded any site that failed to meet the chosen sample size threshold in any analyzed population, and we randomly downsampled when more alleles than the target sample size were available. Populations were excluded for a given arm if the average available sample size was trivially small; this was the case for EA for 2R and 3L, and EG for 2R only. Sample sizes used for X/2R/3L were: CO 9/9/10, EA 14/0/0, EF 44/25/11, EG 21/0/11, FR 60/53/40, KF 12/20/12, RG 22/21/25, SD 43/11/10, ZI 172/150/141. In interpreting data across chromosome arms, we consider (for example), that for arm 2R, the split time of the FR lineage (without EG) from its sub-Saharan ancestral branch refers to the same historical event as the split time of the shared EG/FR lineage for the X chromosome.

Rather than investigate the over two million possible rooted topologies of the above nine populations, we focused on the topology supported by genetic distances (**Figure 3**). We implemented an iterative model-building approach in order to limit the number of simultaneously-estimated parameters. We began with four populations representing the species′ expansion within sub-Saharan Africa: ZI, RG, CO, and EA (or alternately EF). Next, we independently expanded this core model in two directions: (1) by adding the KF and SD samples to complete our southern African sampling, and (2) by adding the EG and FR populations to encompass the species expansion beyond sub-Saharan Africa. For the X chromosome only, we extended the core model a third time to add the EF sample to a model that already contained EA, in order to investigate the timing of the species′ expansion into the Ethiopian highlands. When adding additional populations to our core model, we fixed previous point estimates for parameters already estimated that were not obviously connected to the newly added populations. While this approach might entail minor sacrifices in precision, the ability to include a large number of populations when reconstructing this species’ expansion history amply justifies its use.

Only two branch placements were found to vary among neighbor joining trees^*S4*^ from the three analyzed chromosome arms, and we therefore conducted model choice simulations to evaluate their placement. We evaluated whether the Kafue population should be placed as the basal branch (most likely topology; **Figure 3; Table S4**), or else split from the ZI/SD internal branch, the ZI terminal branch, or the internal branch leading to the non-southern African populations. We also evaluated whether the branch leading to Egypt/France should split from from the terminal branches leading to Rwanda, Cameroon, or Ethiopia, or else from the two internal branches within that population group; of these, a split from the internal branch preceeding the Cameroon/Ethiopia split was most likely (**Figure 3; Table S4**). Model choice was also uses in cases where the temporal sequence of unrelated population splitting events needed to be defined, and again the most likely model was the focus of analysis. For any of these cases, fastsimcoal does not offer a formal test of model choice, and we therefore make no claims about the statistical significance of any given topology. We simply focus on the most likely tree and the demographic estimates associated with it.

Simulations via ms^*S5*^ implementing point estimate models were conducted to confirm agreement with basic summaries of the observed data (Table S5). Confidence intervals for each parameter estimate were obtained by non-parametric bootstrapping - generating 100 bootstrap data sets by resampling with replacement from the empircal data, and performing 100 independent fastsimcoal runs on each of them (mirroring our empirical data analysis). To leverage our estimates from these three chromosome arms, we report the median arm point estimate for each demographic parameter, in addition to arm-specific values. We also report 95% confidence intervals (CIs) for this median arm estimate in addition to CIs for each separate chromosome arm. To generate CIs for the arm median point estimate, one bootstrap parameter estimate was randomly drawn for each chromosome arm, and the middle value was returned.

### Identification of Candidate Regions for Local Adaptation

The *Population Branch Excess* statistic (*PBE^10^*) was used to quantify genetic differentiation specific to the Kafue genomes, when compared against two town-collected populations samples from Zambia (ZI) and South Africa (SP)^*7*^. Due to limited samples of homozygous standard Kafue chromsomes for arms 2L and 3R, inverted chromosome arms were not excluded from the *PBE* analysis, but similar results were obtained from standard arms only.

*PBE* measures the degree to which Kafue-specific allele frequency change exceeds that expected based on locus and genomic differentiation^*10*^. Some loci with strongly positive *PBE* could reflect adaptation to commensal environments shared by the SP and ZI samples, but absent from the Kafue population. *PBE* was applied in diversity-scaled genomic windows containing 250 non-singleton SNPs in the ZI sample. Each window’s *PBE* quantile was determined based on the proportion of windows on the same chromosome arm with an equal or greater *PBE* value.

To assess the genome-wide significance of window *PBE* values, we performed neutral simulations based on an estimated demographic model for these same three populations. For simplicity, we focused on the 2R model which featured an intermediate divergence time between Kafue and the other populations. We typical average window lengths and sample sizes for each population, with a recombination rate of 1 cM/ Mb that should be conservative outside centromeric and telomeric regions^*S3*^. We generated one million ms^*S5*^ coalescent simulations with the following command line: *./*ms 176 1000000 -t 111 -r 169 4600 -I 3 20 134 22 94.2 -en 0 1 0.450 -en 0 2 2.33 -en 0 3 2.18 -em 0 1 3 71.9 -em 0 3 1 71.9 -em 0 2 3 89.4 -em 0 3 2 89.4 -ej 0.00444 3 2 -en 0.00444000001 2 0.542 -em 0.0044000002 1 2 93.8 -em 0.0044000002 2 1 93.8 -en 0.0479 2 0.283 -em 0.0479000001 1 2 4.16 -em 0.0479000001 2 1 4.16 -ej 0.0660 2 1 -en 0.066000001 1 1.01 -em 0.066000002 1 2 0 -em 0.066000002 2 1 0 -en 0.0894 1 1 > KFZISD_test2b.txt

We then obtained a critical value of *PBE* representing a 5% probability that any of 25,548 windows would reach this threshold under the neutral model, under the conservative assumption that each window’s *PBE* value is independent.

The top 1% of *PBE* quantiles were considered outliers for gene ontology (GO) enrichment analysis, under the hypothesis that these outliers will be enriched for genuine targets of local adaptation. GO enrichment was assessed as previously described^4^. Two or more outlier windows were merged into the same outlier window region if they were separated by no more than four non-outlier windows (in order to conservatively avoid counting the same selective sweep more than once). Locations of outlier regions were than randomly permuted, while maintaining their lengths, in order to properly account for the arrangement and lengths of genes in each functional category. Each outlier region was only allowed to vote for a given GO category one time (from both the empirical and permuted outlier regions), in order to avoid spurious results from clusters of functionally linked paralogs.

## DATA AND SOFTWARE AVAILABILITY

Sequence read data is available from NIH Short Read Archive accession PRJNA329555.

## SUPPLEMENTAL ITEM LEGENDS

**Figure S1.** Principle Components Analysis (PCA) indicates a lack of strong genomic differentiation between flies from Kafue National Park (KF) and those from town populations from Zambia (the ZI population sample), South Africa (SA, including the SB, SD, SE, and SF population samples), and Malawi (MW). For each chromosome arm, plots of PC1 (x axis) and PC2 (y axis) are shown. Genomes with detected inversions for a given chromosome arm were excluded in order to search for geographic structure specifically. As expected given the low levels of geographic structure among these populations, each PC explained a small proportion of genetic variance among individuals (noted in parentheses). Occasional non-KF outliers fell outside the regions plotted here. Refers to **Figure 3**.

**Table S1.** Collection information for wildnerness collections cited in the main text, and for two additional wilderness-peripheral collections. Relates to **Figure 1**.

**Table S2.** Individual Kafue genome inversion calls and frequency comparison with Siavonga. Relates to Figure 2.

**Table S3.** Demographic point estimates and confidence intervals. Relates to **Figure 3.**

**Table S4.** Demographic model likelihood comparisons. Relates to **Figure 3.**

**Table S5.** Neutral simulations from demographic point estimates to confirm agreement with genetic variation within populations (nucleotide diversity) and between populations (F_ST_). Relates to **Figure 3.**

**Table S6.** Analysis of *Population Branch Excess* in diversity-scaled genomic windows averaging 4 kb in length. Relates to **Figure 4.**

**Table S7.** Results of gene ontology enrichment analysis, including raw permutation *P* values and the genes within outlier regions for each category. Relates to **Figure 4.**

